# Reconstructing the demographic history of Atlantic Salmon (*Salmo salar*) across its distribution range using Approximate Bayesian Computations

**DOI:** 10.1101/142372

**Authors:** Quentin Rougemont, Louis Bernatchez

## Abstract

Understanding the dual roles of demographic and selective processes in the buildup of population divergence is one of the most challenging tasks in evolutionary biology. In the Northern hemisphere in particular, species genetic makeup has been largely influenced by severe climatic oscillations of the Quaternary Period. Here, we investigated the demographic history of Atlantic Salmon across the entire species range using 2035 anadromous individuals from 77 sampling sites from North America and Eurasia genotyped at 4,656 SNPs. By combining results from admixture graphs, geogenetic maps and an approximate Bayesian computation framework, we validate previous hypotheses pertaining to secondary contact between European and Northern American populations, but also demonstrate that European populations from different glacial refugia have been exchanging alleles in contemporary times. We further identify the major sources of admixture from the southern range of North America to more northern populations along with a strong signal of secondary gene flow between genetic regional groups. We hypothesize that these patterns reflects the spatial redistribution of ancestral variation across the entire American range. Results also point to a role for linked selection in the form of background selection and or positive hitchhiking. Altogether, differential introgression and linked selective effects likely played an underappreciated role in shaping the genomic landscape of species in the Northern hemisphere Therefore we conclude that such heterogeneity among loci should be systematically integrated into demographic inferences of the divergence process, even between incompletely reproductively isolated populations.

## Introduction

An accurate understanding of demographic history, accounting for the putative alternation between periods of isolation and gene flow among populations, is fundamental for population genetic inferences. In particular, the genomic makeup of present day populations in the northern hemisphere is expected to be largely influenced by population splits and secondary contacts linked to climatic oscillations during the last quaternary glaciations (Hewitt 2000). Yet, up to what point the contemporary distribution of genetic variation within species reflects historical divergence processes is a challenging question to address. Under the genic view of speciation, during allopatric phases, populations can randomly accumulate genetic Dobhzansky-Muller incompatibilities and other genetic barriers to gene flow due to genetic drift and selection (Wu 2001; Harrison and Larson 2016). Following secondary contact, gene flow is expected to erode past genetic differentiation outside of barriers. Depending on the balance between levels of secondary gene flow and number of accumulated barriers in allopatry, heterogeneous landscapes of divergence at the genetic level may arise (Wu 2001; Wolf and Ellegren 2016). In line with this expectation, empirical population genomics studies from the last few years have documented the near ubiquity of the heterogeneous landscape of differentiation across a continuum of increasing divergence (reviewed in Seehausen et al. 2014; Wolf and Ellegren 2016).

However, the observation of these 'islands of divergence' have fueled the long standing debate about whether populations and species are currently diverging in the face of continuous gene-flow or through allopatric divergence, emphasizing the need to reconstruct the initial conditions of population divergence and subsequent gene flow. Earlier studies have used genome scans to identify islands of differentiation (Feder et al. 2012; Seehausen et al. 2014) and draw verbal theory about ‘speciation islands’ as a basis for divergence with gene flow. Under a model of divergence hitchhiking, gene flow keeps eroding genetic differentiation outside of selected regions while selective sweeps involved in local adaptation and linked neutral variants will show higher level of differentiation (Via and West 2008; Feder et al. 2012; Flaxman et al. 2013). As stated above, however, allopatric divergence followed by secondary contact, whereby gene flow will erode past genetic differentiation outside of barrier loci or where gene flow is less effective (Barton and Bengtsson 1986), can produce the same patterns (e.g. Gagnaire et al. 2013; Burri et al. 2015; Rougeux et al. 2016; Roux et al. 2016; Rougemont et al. 2017). Moreover the link between heterogeneity of differentiation and the role of gene flow during divergence is increasingly questioned as hitchhiking of neutral alleles linked to a selective sweep, especially in regions of low recombination, can produce similar pattern (Noor and Bennett 2009; Cruickshank and Hahn 2014). Similarly, background selection (Charlesworth et al. 1993; Charlesworth 1994) can contribute to this heterogeneous genomic landscape of divergence. This linked selection can be seen as an increase in genetic drift and thus be modeled as a local reduction of effective population size (*Ne*). This was first demonstrated by Hill and Robertson (Hill and Robertson 1966) who proposed that interference among loci under selection can be approximated as an increase in genetic drift, as compared to the effect of selection on single loci. Under these linked selective process, *Ne* will be reduced in genomic areas of low recombination relatively to regions of higher recombination (Hill and Robertson 1966; Charlesworth et al. 1993). Neglecting such selective effects in demographic inference can lead to biased inferences (Ewing and Jensen 2016; Schrider et al. 2016). Therefore, a modeling approach that jointly allows for local genomic variations in effective population size and migration rate can improve our understanding of the demographic processes at play during population divergence. Simulation tools such as approximate Bayesian computations (Tavaré et al. 1997; Beaumont et al. 2002) allow for the simulation of complex models of divergence history. New methods have been developed to model the various levels of introgression across the genome (Roux et al. 2013; Sousa et al. 2013; Tine et al. 2014), as well as local variation in effective population size (Tine et al. 2014). Altogether, these approaches may help understanding the joint effects of barrier loci reducing migration rate and of linked selection reducing local effective population size.

Atlantic Salmon (*Salmo salar*) is a particularly relevant model to study the interactions of contemporary and historical factors leading to heterogeneous landscapes of divergence. Distributed throughout the North Atlantic, both in Eastern North America and Europe, it undergoes long anadromous migrations to feed at sea at the adult stage before returning to natal rivers for spawning (Quinn 1993). Such homing behavior results in reduced gene flow at both local and regional scales that in turn translate into fine scale spatial structure and a pattern of isolation by distance, which altogether may also facilitate the establishment of local adaptation (Taylor 1991; Dionne et al. 2008; Perrier et al. 2011; Primmer 2011). Accordingly, a large body of literature has been devoted to describing the population genetic structure of the species (Vasemägi et al. 2005; Palstra et al. 2007; Perrier et al. 2011; Bourret et al. 2013; Moore et al. 2014), documenting local population sizes (Palstra et al. 2009; Ferchaud et al. 2016) and the effect of hatcheries on population admixture (Milot et al. 2013; Perrier et al. 2013; Bolstad et al. 2017). In contrast, far less is known about the demographic history of the species, globally, as well as within each continent. Previous studies have documented pronounced continental divergence between European populations and Northern American populations using various molecular markers (Cutler et al. 1991; Bourke 1997; King et al. 2007; Bourret et al. 2013). It has been proposed that continental divergence most likely occurred around 600,000 −700,000 years ago (King et al. 2007) and perhaps more than one million year ago (Nilsson et al. 2001). The process resulted in karyotypic divergence between North American and European populations, possibly due to Robertsonian fissions and fusions (Hartley 1987) as well as to other types of chromosomal rearrangements (Lien et al. 2016). Yet, European alleles from mtDNA and allozymes have been found to segregate at low frequency among American populations from Newfoundland and Labrador (Verspoor et al. 2005; King et al. 2007). In contrast with this deep continental phylogenetic divergence, European populations likely split more recently and following the end of following the Last Glacial Maxima (LGM) around 18,000 years ago, expanded from potential southern refugia in the Iberian peninsula and other non-glaciated rivers further north (King et al. 2007). Recent studies (King et al. 2007; Bourret et al. 2013; Bradbury et al. 2015) have also supported such a historical scenario whereby salmon populations may have diverged in multiple refugia within Europe, and whether these populations have come into secondary contact has not been resolved yet. In contrast to the pronounced regional level of population genetic structure among European populations, the North American populations show a smaller amount of genetic diversity and lower level of genetic differentiation (King et al. 2007; Bourret et al. 2013). What is missing towards further elucidating the origin of contemporary population structure in the species is a thorough investigation of the role of different historical factors, which would explicitly take into account the potential effects of linked selection.

The goal of this study was to reconstruct the Atlantic Salmon demographic and population divergence history between the two continents as well as within Europe and North America. To do so, we took advantage of two previously published dataset that have solely focused on describing patterns of population genetic structure (Bourret et al. 2013; Moore et al. 2014) but did not perform explicit demographic inferences. First, we used classical population genetic clustering and tree-based approaches to identify the most likely source of divergence. Second, we used an accurate Approximate Bayesian Computation (ABC) framework and compare alternative models of population divergence that integrate both genome-wide variation in effective population size and migration rate in order to jointly account for the effect of linked selection and barriers to gene flow.

## Results

### Population genome-wide diversity and divergence

After excluding 45 individuals with more than 5% missing genotypes as well as SNPs with a lower than 95% genotyping rate a total of 2,035 individuals from 77 sampling locations (Fig. 1a, Supplemental Table S1) with 4,656 SNPs were kept for all subsequent analyses. A significantly lower observed heterozygosity among Northern American populations (Ho = 0.16) compared to that observed in Europe (Ho = 0.29; p-value <0.0001; Supplemental Table S2) was observed. Global differentiation was largely heterogeneous across the genome with a mean *F*_ST_ value of 0.349 across all populations (min = 0.033, max = 0.971, Supplemental Fig. S1). Mean *F*_ST_ among European populations was lower (*F*_ST_ = 0.142 min genome-wide = 0.00 – max = 0.68) than that observed between continents and level of differentiation among Northern American populations was lower than that observed in Europe (mean *F*_ST_ = 0.090, min genome-wide = 0 − max = 1), see also Supplemental Table S2 and S3). Based on an analysis of molecular variance stratified at the continental level, 46 % of variance was observed between continents (p > 0.001), 47 % of variance was observed within population and 7 % of variance was observed among populations within continents.

**Fig. 1:**
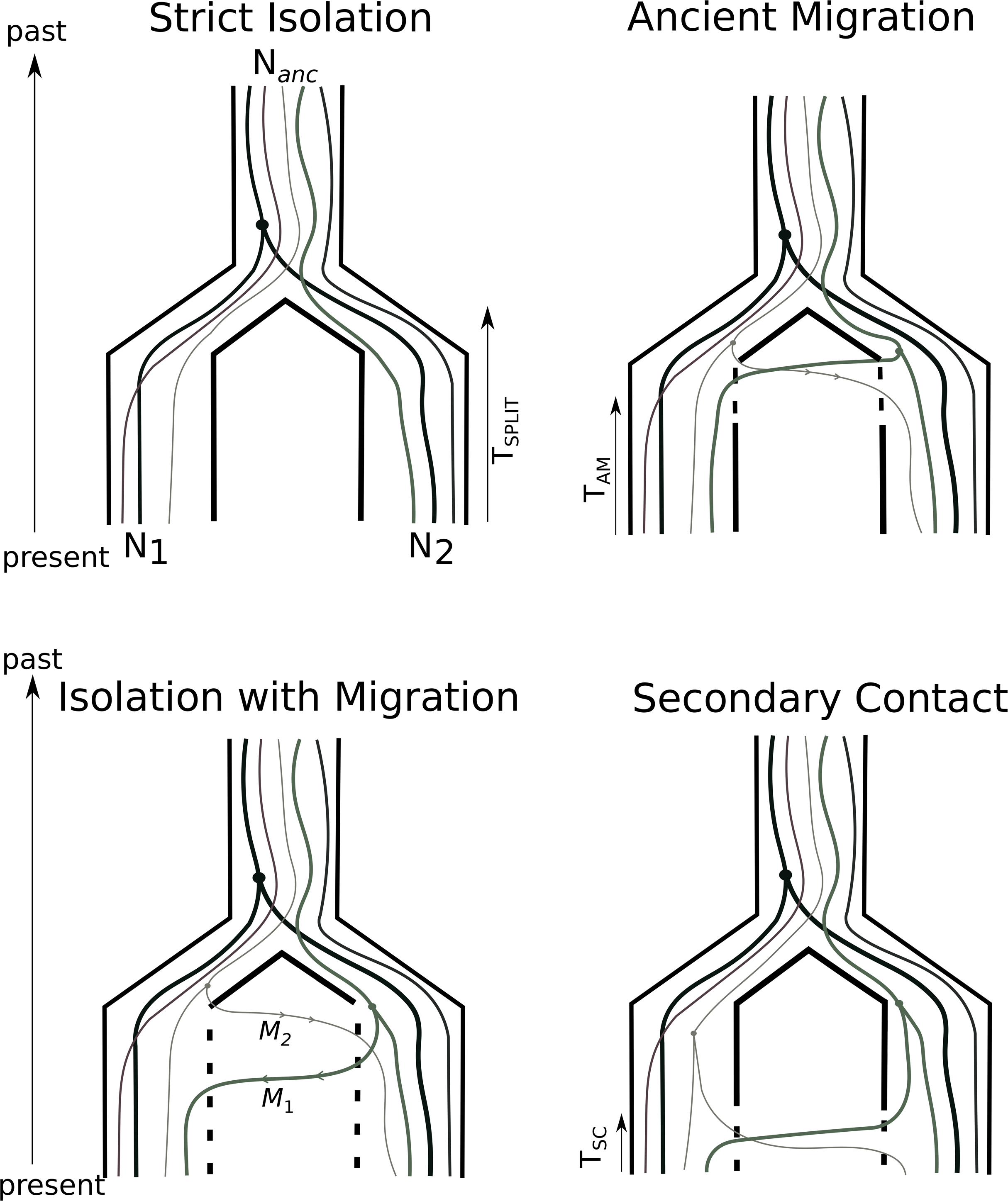
Representation of the demographic scenarios compared in this study: Strict Isolation (SI), Isolation with constant Migration (IM) Ancient Migration (AM) and Secondary Contact (SC). All models share the following parameters: *T_split_*: number of generation of divergence (backwards in time). Nanc, *N_1_*, *N_2_*: effective population size of the ancestral population, of the first and second daughter population compared. M_1_ and M_2_ represent the effective migration rates per generation with *m* the proportion of population made of migrants from the other populations. *T_am_* is the number of generations since the two populations have diverged without gene flow. *T_sc_* is the number of generations since the populations have started exchanging gene flow (secondary contact) after a period of isolation.

### Individual clustering

We explored the broad patterns of population genetic structure using principal component analysis (PCA). The first axis of the PCA (Fig. 2a and Fig. 2b) captured the majority of the differentiation with 33.84% of the total inertia, and clearly separated populations from each continent, consistent with the strong mean inter-continental *F*_ST_ value. All North American populations clustered together on the first axis while the second axis (2.31% of inertia) separated European populations into three major groups corresponding to an “Atlantic” group, a group from the Northern Europe (Barents and White sea) and a Baltic group as defined previously (Bourret et al. 2013). Plots of the axis 1 and 3 and axis 1 and 4 were very similar to the first and second axis as most differentiation was driven by continental divergence (Supplemental Fig. S2). Projection of the axis 2 and 3 of the PCA captured 4% of inertia; (Fig. 2b) and separated mostly the Baltic Sea populations from all others while North American populations fell at the center of the axes and the Atlantic populations clustered along the first axis with a tendency toward a North-South organization. Population from the Barents-White Sea clustered separately along the second axis and here, westernmost (e.g.Tuloma, Tana) populations were closer to North American populations while easternmost (e.g. Suma, Pongoma) populations were further apart along this second axis.

**Fig. 2:**
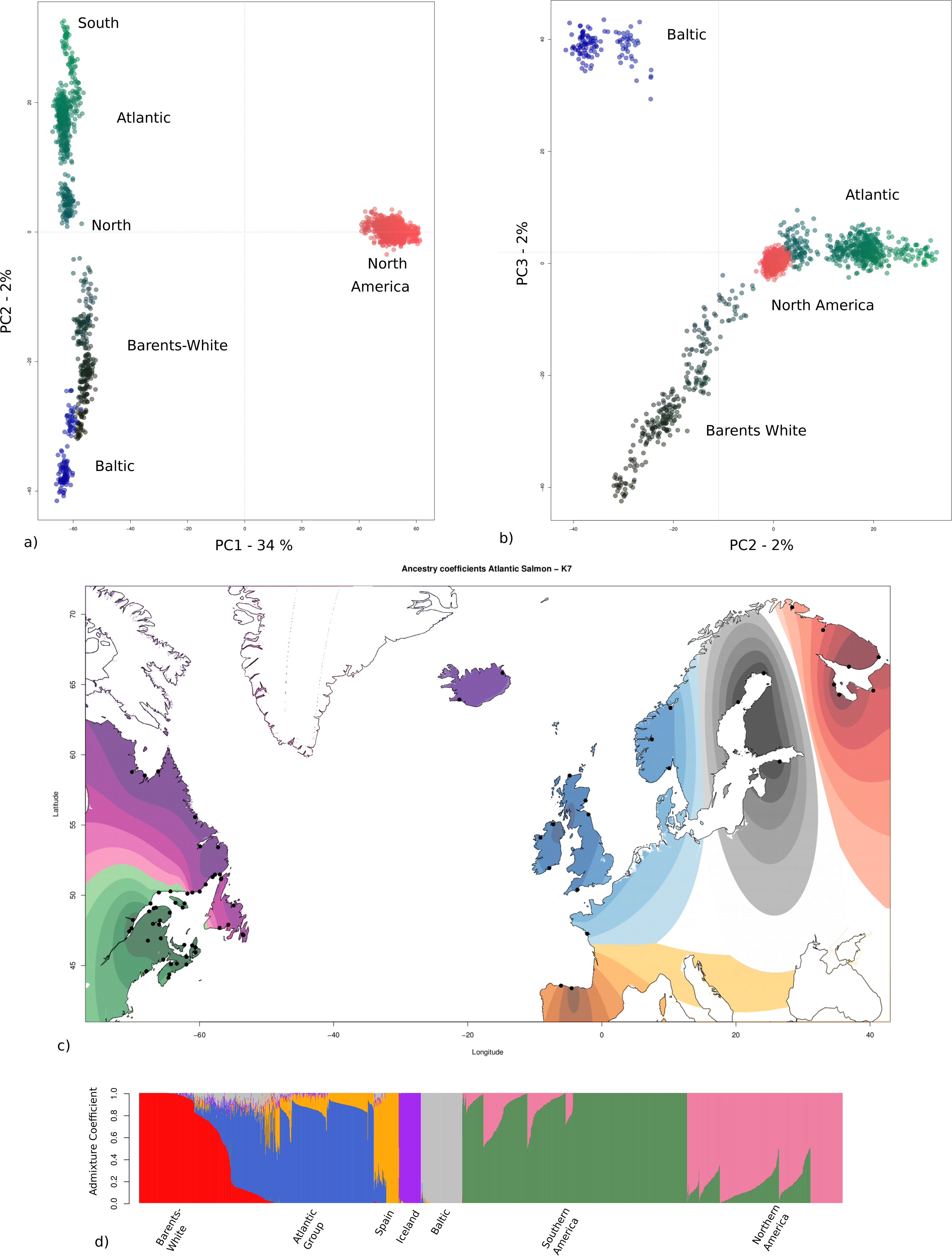
Patterns of population genetic structure among all 2035 Atlantic Salmon analysed in this study. a) Principal components analysis with the first two axes plotted; b) PCA of the axis 2 (horizontal) and 3 (vertical); c) Spatial interpolation of population structure inferred for K = 7 clusters. Each black dot represents a sample points. Colors depicts ancestry coefficient of the major groups. Spatial interpolation of ancestry coefficient was obtained using a kriging method following Jay et al. (2012). We mapped only cluster with the maximum local contribution to the observed ancestry on each point. Interpolation outside of the species range is not relevant for interpretation. The associated barplot is provided in d) where each vertical line represents an individual and each color represents a genetic cluster. Individuals are sorted according to their membership probability. Major geographic areas are provided below the plot. Results for additional K-values are provided in Supplemental Fig S4 and S5.

Results of population structure using the R package LEA (Frichot and François 2015) suggested that estimating a single fixed K was difficult (Supplemental Fig. S3) as the crossentropy criterion softly decreases and indicated that several clustering values could fit well the data. We therefore plotted a diversity of solutions to avoid over-interpretation (Falush et al. 2016). A first change in the minimal cross-entropy criterion was observed for K =5 and K= 7 (Supplemental Fig. S3). K = 5 revealed the same clustering as the PCA (Supplemental Fig S4). At K = 7 (Fig. 2c and barplot Fig. 2d), 42% of individuals showed a q-value <0.9 value (Fig. 2d). The seven groups inferred corresponded to the 'Baltic' group (mean q-value = 0.99), the ‘Icelandic’ group (mean q-value = 0.99), the 'Spanish group' (mean q-value = 0.97), the 'Atlantic' group (mean q-value = 0.82), the 'Barents-White sea' group (mean q-value = 0.86) a “Southern” group in America (mean q-value = 0.82) and a Northern Group in America (mean q-value = 0.85). Individuals from the southern parts of the Barents-White Sea group displayed an ancestry coefficient closer to that of the individuals from the Atlantic group, forming a cline with those individuals. Similarly, the Loire River population (Atlantic group) displayed consistently mixed membership as no salmon displayed a q-value greater than 0.90 and shared ancestry with salmon from Spain. Moreover, 36 individuals from two Spanish rivers displayed mixed ancestry with individuals of the Atlantic group while the remaining individuals from a single Spanish river were assigned to this 'Spanish' group with a q-value >0.99. In contrast, individuals from the Baltic Sea and Iceland were all assigned to these 'regional' groups with a q-value greater than 0.90. Salmon from North America were separated into a Northern and a Southern group with respectively 194 out of 736 (26%) and 156 out of 359 individuals (43%) displaying mixed membership. The boundary delimiting the Northern and Southern population was falling in the Natashquan River from the Lower Shore of Québec.

Investigating a K value of 13 (Supplemental Fig S4)where the cross-entropy criterion further dropped separated individuals from Europe into eight subgroups, with notably the Loire and rivers from the Spanish area (Cares, Narcea, Piguena) that clustered separately as well as the most geographically isolated populations within the Baltic and Barents areas. Salmon from North America were further separated into five major groups exhibiting variable levels of admixture but with a tendency towards a geographical clustering according to a north-south gradient. Lastly, investigating higher levels of clustering (K values of 20, 30 or 40, Supplemental Fig. S5) revealed increasingly higher levels of population admixture within North American groups, with mean q-value, (averaged for all cluster) ranging from q = 0.75, q = 0.62 and q = 0.65 for K = 20, 30 and 40 respectively. In contrast, levels of admixture among differentiated clusters in Europe were less pronounced as q-value were higher (mean q-value = 0.90, 0.88 and 0.87) for K = 20, 30 and 40 groups respectively (Supplemental Fig. S5). Overall, our results corroborate those of two previous studies (Bourret et al. 2013; Moore et al. 2014) but also allowed better defining major genetic groups in some cases, mainly in Europe.

### American population were founded by multiple European sources

We combined TreeMix v1.12 (Pickrell and Pritchard 2012), *f3-test* (Reich et al. 2009) and Spacemix (Bradburd et al. 2016) to identify population splits and mixture as well as the most likely source or sources of admixture. *Treemix* analyses were performed using regional genetic groups defined in Bourret et al. (2013); Moore et al. (2014) and complemented by our own clustering analysis (Supplemental Table S1). This pooling strategy further allows to reduce the noise due to ongoing gene flow between neighboring rivers and to simplify the visualization of the tree in order to test the major migration events. A total of 4149 SNPs that were successfully placed on the Atlantic Salmon reference genome (Lien et al. 2016) was used.

*Treemix* indicated a steady increase in the percentage of variance of the covariance matrix explained as migration events (red arrow in Fig. 3b) were added to the tree (Supplemental Fig. S6). Indeed, 99.8% of variance in the covariance matrix was already explained without migration. However, this percentage increases for different number of migration events and up to 19, demonstrating that even if most of the history of divergence can be summarized without gene flow, a more complex history of split and subsequent gene flow provided better fit to the data. We compared the results without migration (Fig. 3a) and with nine migration events (Fig. 3b). Results for values of *m = 1* and *m* = 3 migration events are provided in Supplemental Fig. S7 together with plots of the residuals in Supplemental Fig. S7 and S8. Considering between one and four migrations events revealed that the most important migration events consistently occurred from the Baltic Sea to the Barents Sea in Europe and from Nova Scotia (NVS) to Avalon in North America. For nine migrations events, admixture occurred again from the southernmost American populations (NVS and Narraguagus (Nar, Maine)) to three other regional groups (Avalon, Newfoundland and Labrador). We also detected admixture from European populations to Gaspésie and Anticosti regions in North America. Lastly, there was evidence for three admixture events along the branch leading to the Baltic group into one ancestral branch leading to the Barents-White sea group and in two other branches of the Barents-White sea population (Tana-Tuloma Rivers and Emtsa River).

**Fig. 3:**
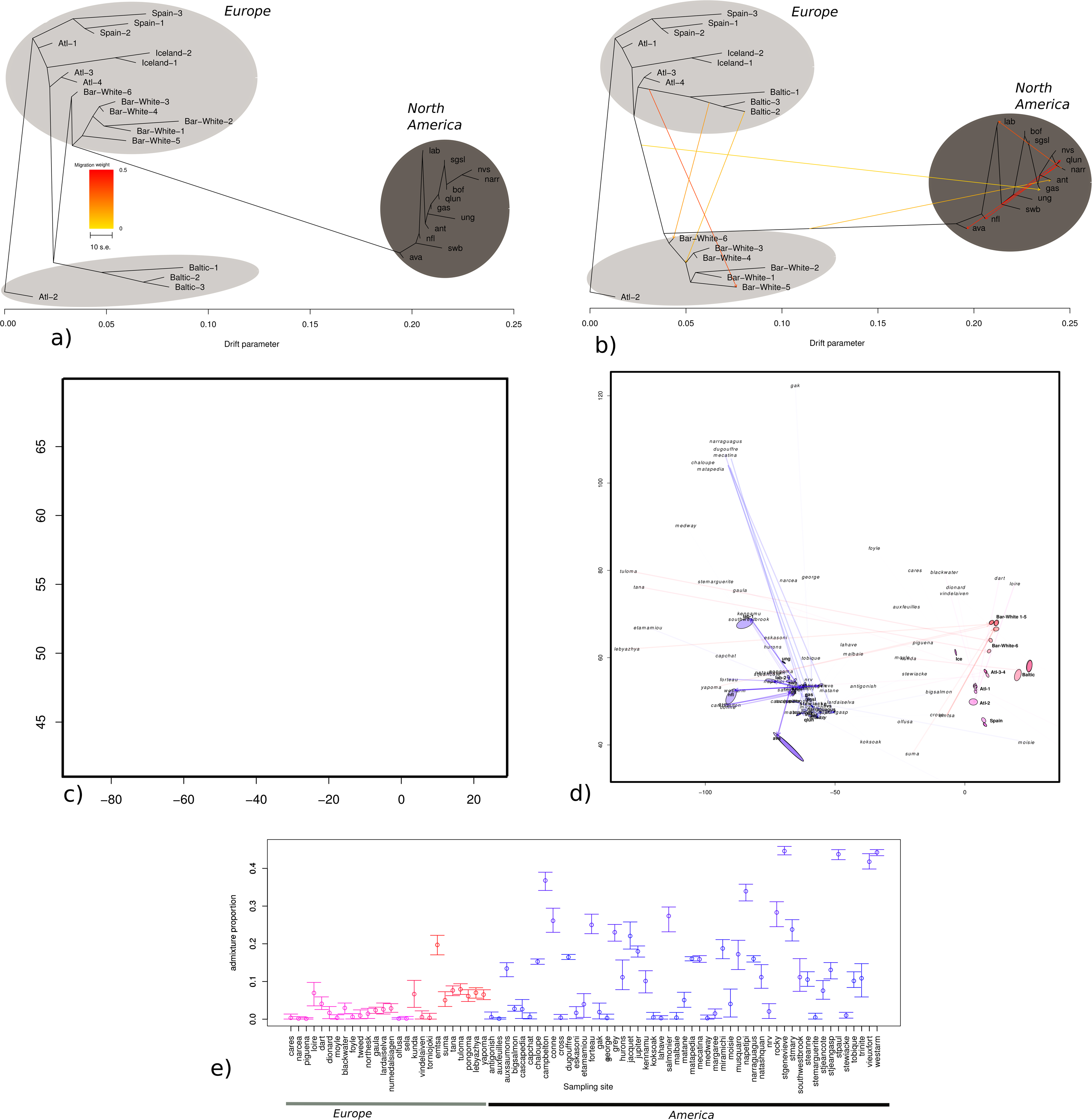
Population admixture and sources of gene flow. a) Admixture graph inferred with Treemix without migration and b) *m* = 9 migration events. The x-axis represents the amount of genetic drift, proportional to *Ne*. Arrows depict admixture events and color gradients indicate the strength of admixture. Details of population pooling are provided in S1 Table. c) Spacemix map inferred, without admixture corresponding to a model of pure isolation by distance. Each name represents a sample site and the color gradient is made according to each sample geographic position. The map broadly recovers the results of the PCA with American sample clustered on the left side (blue gradients) and European samples clustered on the right side (red gradients). d) Spacemix inferred geogenetic map with admixture. Source of admixture are represented in italics and the original position of each sample points are in bold. For America, original positions were removed for ease of interpretations and replaced by the “major groups” from which they belong (S1 Table). Arrows joins the location of admixture source (in italics) to the current location of the sample, opacity of the arrow is a function of the amount of admixture of the sample. Ellipses around each point represent the 95% CI of geogenetic locations. e) Admixture proportions inferred by Spacemix for each sampling locations with 95% CIs. Color scheme is the same as before with color gradient based upon geographic position.

For each triplet of populations, the *f3*-statistic was used to formally test if one population draws significant admixture from the two other populations. Out of 447 statistically significant tests, eight regional genetic groups from North America and one from the Barents Sea (Tuloma River) appeared significantly admixed with genetic background from other such groups (Supplemental Table S5). The Baltic Sea was a source of admixture towards the Barents Sea in 80% of the significant tests, confirming Treemix results for *m* between 1 and 5. In North America, the most statistically significant sources of admixture were the two southernmost American groups of Nova Scotia (NVS, 150) and Narraguagus (Nar, 150). Importantly, significant *f3*-tests of admixture into American populations always involved one population of Europe with one American population, whereas not a single test involving two European populations or two North American populations was significant. Therefore, our *f3*-tests provide evidence for the hypothesis that American populations resulted from admixture from several historical sources of American ancestry (here associated with Nova Scotia and Narraguagus) with most European ancestral populations.

To get further insights into the levels of admixture among genetic groups, Spacemix was used to construct ‘geogenetic’ maps where the distances are a function of the level of genetic differentiation between populations evolving under a model of Isolation-by-Distance. The occurrence of admixture then resulted in a lower than expected geogenetic distance. Since this analysis allows incorporating geographic information, it was performed at the level of sampling sites, without grouping into higher order groups as in the previous analyses. Results show that geogenetic maps without admixture largely recover results expected from a simple PCA (Fig. 2a *versus* Fig. 3c), however a model incorporating both admixture and migration provided additional information regarding the demographic history of populations (Fig. 3d). It is complicated to provide a detailed description of all admixture events, as all populations displayed admixture and occupy a different geogenetic position. Nevertheless the visualization of the geogenetic map indicates that four European populations (Tuloma, Tana, Lebyazhya and Lardaiselva have their admixture source close to American populations. Similarly, the Moisie, Cross, Emtsa, AuxFeuilles and Stweiacke Rivers from North America have admixture source closely related to European populations. Moreover, another five populations (Chaloupe, DuGouffre, Narraguagus, Matapedia, Mecatina) have their admixture source closely positioned on the map and are distant from the actual geogenetic position of the sample. The same applies to the Gaspereaux River with its admixture source being located at the top of the map.

In agreement with this result and the *Treemix* and PCA analyses, Fig. 3e shows that most North American populations display higher levels of admixture than European populations with a mean admixture proportion for North America of 0.136 [95%CI=0.117−0.156] versus 0.036 [95%CI=0.024-0.051], which were significantly different (Wilcox test W = 350, p-value < 0.005). In addition to this significant difference, it is noteworthy that the variance was very large with some American populations displaying low admixture levels similar to those observed in Europe (e.g. Cross, Koksoak, Malbaie Fig. 3e) while others displayed high admixture proportion (eg. St Paul, Westram, Fig. 3e). Finally, an analysis at the “regional” level, using the same groups as in *Treemix* largely recovered the results with the Narraguagus, Bay of Fundy; St-Lawrence and Avalon group having their source of admixture placed close to the European Atlantic group (Supplemental Fig. S9).

### Demographic history and divergence

While all the above analyses were informative with regards to the most probable source of ancestral admixture, their underlying demographic models are simplified and it is not possible to explicitly fit a model of population divergence. Therefore we performed an explicit modeling approach testing the four different models (Strict Isolation, Isolation with Migration, Ancient Migration and Secondary Contacts) including the possibility for heterogeneous introgression rate (*Mhetero*) and drift (*Nhetero*) among loci, as outlined in the Methods section. Using these models, we performed a total of 163 comparisons between pairs of populations. Demographic inference were performed by pairs for the following reasons: *i)* classic scenario of divergence are general compared between pairs of population *ii)* the number of parameters involved in a three or four populations models including heterogeneous migration and effective population size becomes too high and would certainly requires a higher amount of data than that available here. Thus a total of 90 comparisons were performed between the European and American continents, 37 comparisons between American population pairs from different major regional areas identified previously (Moore et al. 2014), and 36 comparisons were performed between European population pairs from the major regional groups proposed in another study (Bourret et al. 2013) and also corroborated here.

### Model selection and robustness

The most striking pattern emerging from the model selection procedure was the high posterior probability of the secondary contact model when averaged over the four declinations (*NhomoMhomo, NheteroMhetero, NheteroMhomo, NhomoMhetero*). The median averaged over all comparisons was P(SC) = 0.99 (sd ± 0.091) (Fig. 4, Supplemental Table S6) whereas 88% of the comparisons had a P (SC)>0.90 with a robustness greater than 0.91 as measured using 140,000 PODS (i.e.14 models × 10,000 PODS). Eighty-four percent of the comparisons displayed robustness greater than 0.95 and a posterior probability of SC > 0.969. Also, comparisons between Homo model (*NhomoMhomo*) versus Hetero models (*NheteroMhetero, NheteroMhomo, NhomoMhetero*) showed that models incorporating heterogeneity of either *Ne, m*, or of both were largely favored, as indicated by a higher posterior probability (Fig. 5). For a random subset of 32 out of the 148 models with robustness above 0.865, we tested whether integrating two episodes of isolation and secondary contact (periodic secondary contact, PSC, see details in methods and Supplemental Note S1) or whether including postdivergence bottlenecks (BSC) improved the fit. In the 32 cases tested, the ABC model selection indicated that there was little gain from these models (Supplemental Table S7). Thus, we based our inference on their least parameterized alternatives. To ascertain the potential effect of contemporary admixture, due for instance to recent stocking of Atlantic salmon from genetically differentiated rivers, we run another set of abc analysis (*n* = 20) on a random subset of individuals assigned to their genetic sampling site with a q-value > 0.90. The P(SC) averaged over all comparisons was > 0.93 (Supplemental Table S8). Therefore, we conclude that recent admixture due to human activity should have minimal effects on inferences using all individuals and only present our results based on the whole set of individuals in all pairs.

**Fig. 4:**
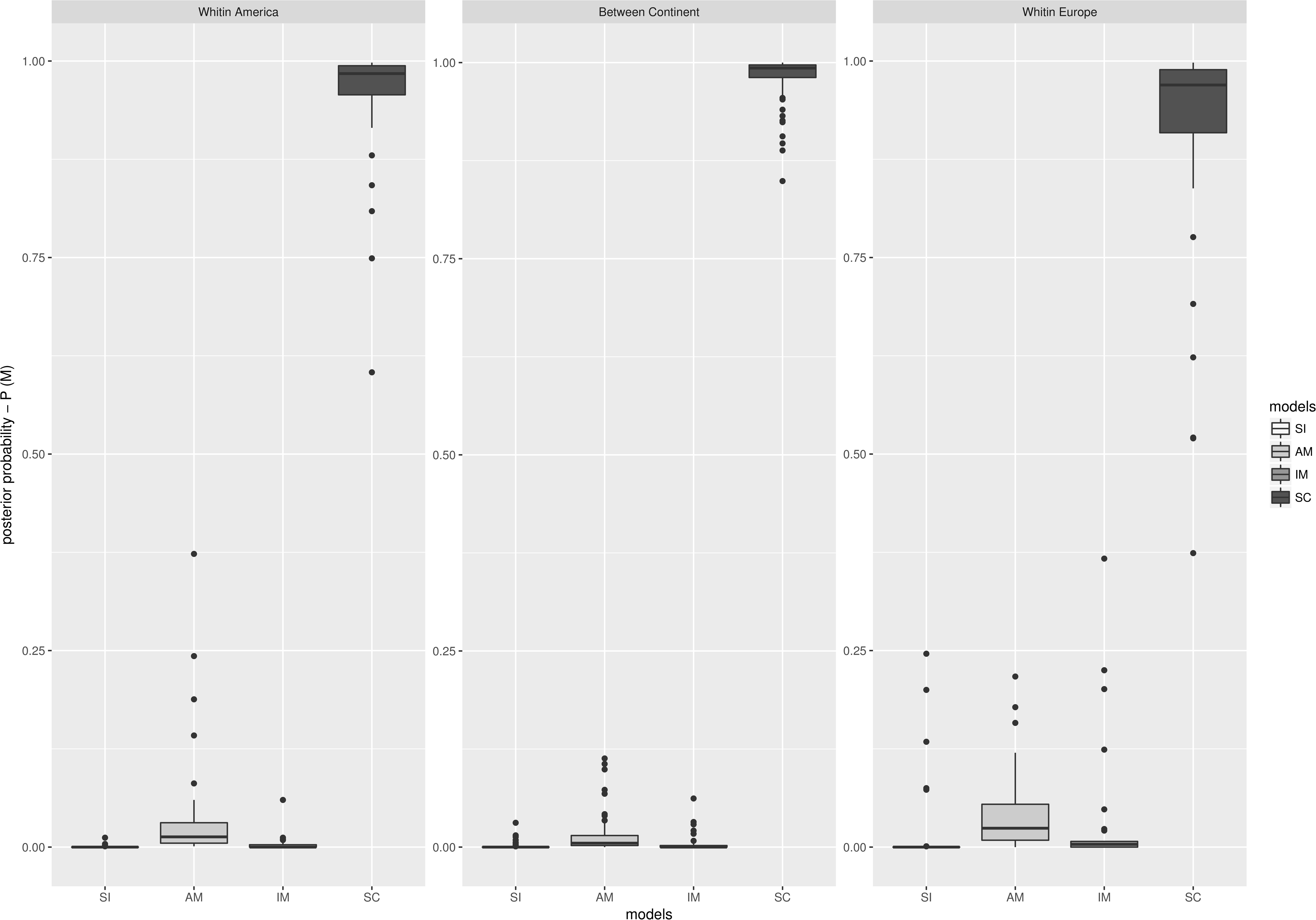
Boxplot of the posterior probability of each of the four compared models. Each boxplot is composed of the posterior probability obtained in “between continents” comparison *n =90*, “within America” comparison *n = 37*, and “within Europe” comparison *n= 36*. SI= Strict Isolation, IM = Isolation with Migration AM= Ancient Migration, SC = Secondary contact. The 14 models were compared at the same time and the posterior probability of each possible version of SI, AM, IM and SC was summed.

**Fig. 5:**
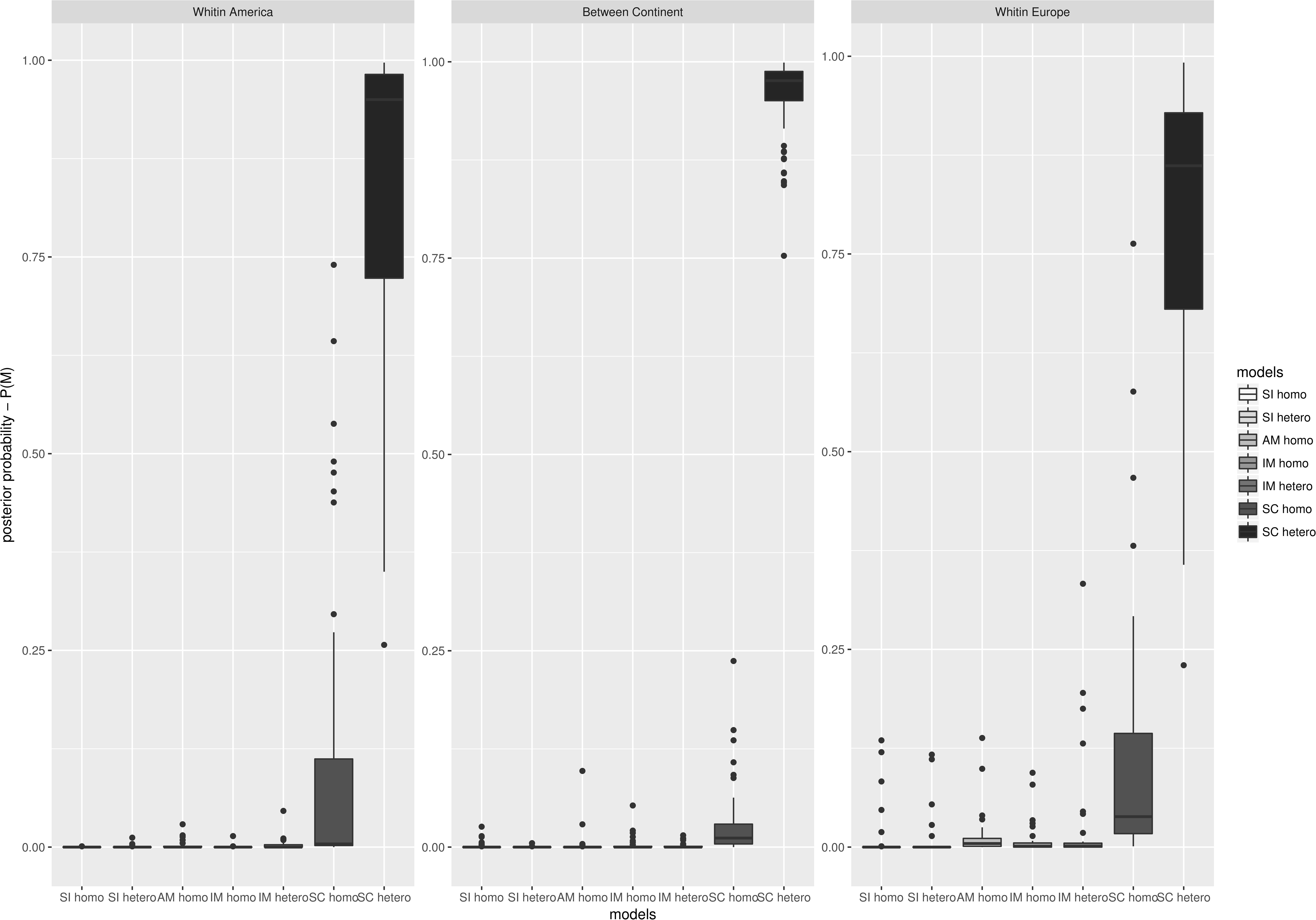
Posterior probabilities of homogeneous versus heterogeneous models. Each boxplot is composed of the posterior probability obtained in “between continent” comparison *n =90*, “within America” comparison *n = 37*, and “within Europe” comparison *n= 36*. The posterior probability of each of the four models incorporating either heterogeneous introgression rate (*m*), heterogeneous population size (*Ne*) or both (*m* and *Ne*) was sum against model assuming homogeneous introgression rate and population size.SI= Strict Isolation, IM = Isolation with Migration AM= Ancient Migration, SC = Secondary contact

### Parameter estimations and goodness of fit

Demographic parameters were estimated for a subset of models (n = 148) with robust inferences at a threshold posterior probabilities ≥ 0.865. We first performed a posterior predictive check to test the accuracy of our simulation procedure. We found that under each of the best model, the simulation pipeline produced accurate estimates based on the posterior distribution of 2,000 newly simulated summary statistics. Of the 148 comparisons, we found that π tended to be significantly different from the observed value (p < 0.001) in a total of 46 cases. The average and standard deviation of the number of fixed differences (Sf) was also less accurately reproduced with the simulation pipeline in a few cases (n = 16, p < 0.001) (Supplemental Table S9). Parameter estimates were then performed under the best scenario of secondary contact among *NhomoMhomo, NheteroMhetero, NheteroMhomo, NhomoMhetero*.

Posterior parameter estimates were well differentiated from the prior for the time of secondary contact in 147 out of the 148 comparisons and indicated very recent secondary contacts as they represented on average 1.3% of the split time between continent, 1.5 % of the split time between population within America Europe and 0.7% between populations within Europe (Fig. 6). In contrast, posterior estimates of split times were differentiated from the priors in 13% of the comparisons, and all but two were between populations from North America (Supplemental Table S10). Given the importance of estimating split time but the high uncertainty surrounding its estimation, the following complementary approach was used. We identified for the 148 most robust comparisons the most differentiated loci according to their *F*_ST_ values. All loci falling outside the 95% percentile of the *F*_ST_ distribution were sampled and parameters were estimated under a model of strict isolation SI. For a few cases where all loci were fixed at the 95% level we used a 90% threshold. Under the genic view of speciation (Wu 2001), barrier loci harboring various genetic incompatibilities are more likely to have been resistant to introgression, and to display deeper coalescent times, therefore having high *F*_ST_ values. In these conditions, a less parameterized model of strict isolation appears as a parsimonious alternative to estimate split time. The whole simulation pipeline was used as above (i.e. one million simulations for 148 pairs) using this restricted set of markers. A model selection choice was performed in a subset of the data (n = 60) in order to ensure that the SI model was indeed the best model. Between continents, we found that at a 95% *F*_ST_ percentile threshold, the SI model was indeed the best model except in 14 cases generally involving the same river (Yapoma River). However, this was not the case for within-continent comparisons, where the SC always remained the best model. The 90% threshold produced similar results but with the discrimination between SC and SI being less clear. For within continent comparisons, we found that posterior estimates of split times under SC were systematically not differentiated from the prior except in four cases and consequently not interpretable. We therefore present only estimates of split time under SI between continents. In this configuration, divergence times were accurately estimated in 65% of the comparisons (Supplemental Table S11). Our inference indicated a mean continental split time of 20.84 coalescent units (Supplemental Fig. S10), which was significantly higher than the mean split time of 12.61 coalescent units inferred from all markers (*t*-test, p <0.0001). It is noteworthy that in the four estimates under the Secondary Contact model at the 95% threshold where the posteriors were well differentiated from the prior, the split time was also older than using all markers (mean = 26.44 coalescent units), reflecting the possibility that at the genomic level, accumulation of genetic barriers involved in adaptation and/or isolation predated the onset of divergence.

**Fig. 6:**
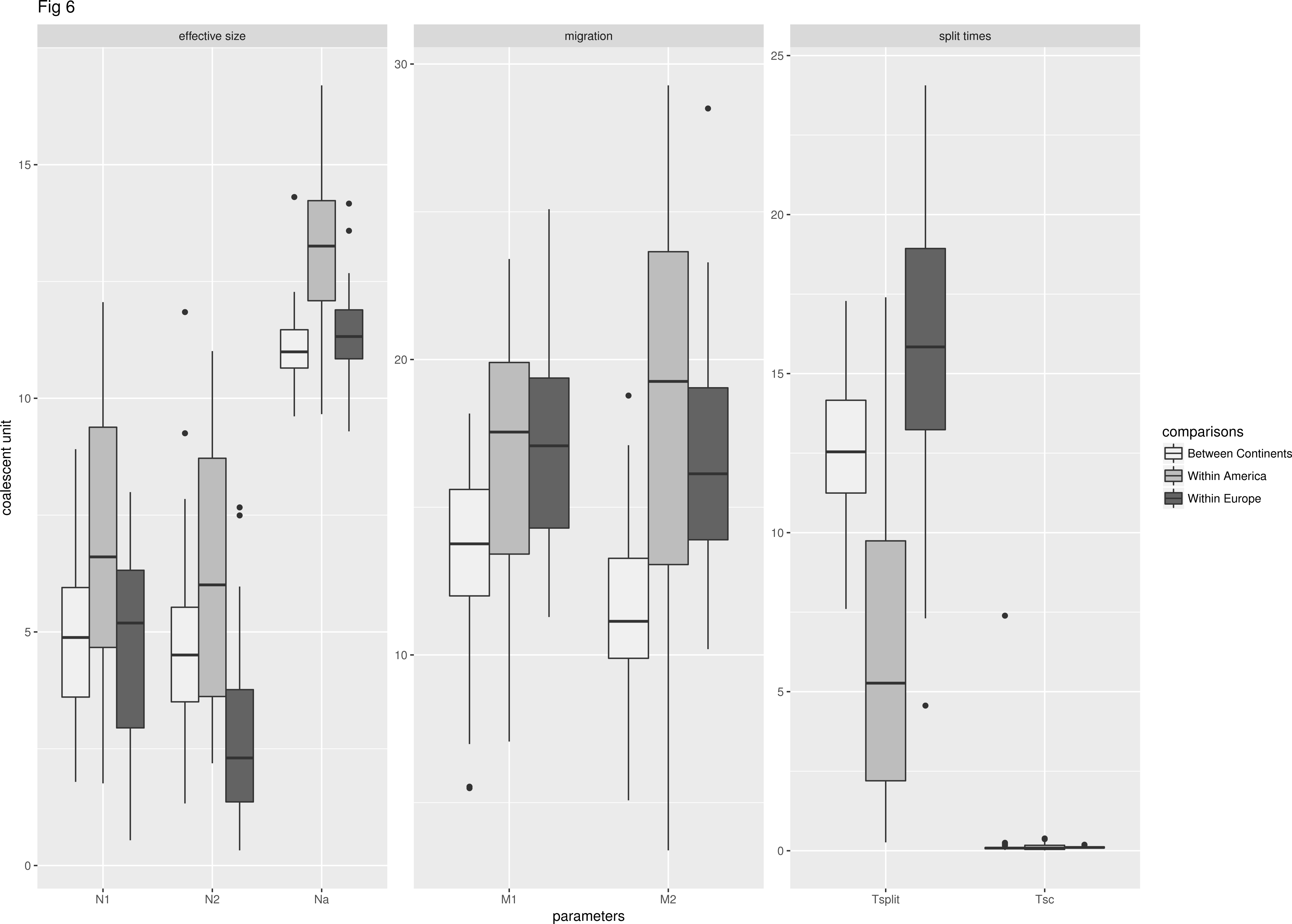
Estimates of demographic parameters under the best secondary contact models. Mean values are provided and averaged over all comparison between continent, within American populations and within European populations.

Posterior estimates under the secondary contact model also indicated a reduction in effective population size of the contemporary populations compared to the ancestral population size but with large confidence intervals. This result was confirmed by estimates under the strict isolation model where the differences were significant (*t*-test p < 0.0001). Under the SI model, we also found that North American populations displayed significantly higher contemporary population sizes than European populations (*t*-test p < 0.0001; Supplemental Table S10). Estimates of migration rate under the secondary contact models produced large confidence intervals, with posterior being differentiated from the prior in less than 50% of the comparisons making the interpretation of those results not relevant (Supplemental Table S10).

## Discussion

The main goal of this study was to reconstruct the demographic history of between-continent and within-continent divergence of Atlantic Salmon populations across the northern hemisphere. Towards this end, an ABC framework was implemented, taking into account the effect of linked selection locally reducing *Ne*, and of genetic barriers, locally reducing the rate of effective gene flow (*m*). Demographic inferences strongly supported a long period of geographic isolation, during which linked selection likely shaped the heterogeneous landscape of divergence between them. Periods of isolation were subsequently followed by widespread secondary contacts between continents as well as within each continent which have contributed to the erosion of genome-wide differentiation outside of genetic barriers.

### Population structure

The PCA and clustering analyses largely recovered results of two previous studies (Bourret et al. 2013; Moore et al. 2014) for North American and European populations and confirmed that *i)* the largest divergence occurred between European and North-American populations, with 36% of the variance attributed to between continents divergence; and *ii*)pronounced regional genetic clustering occurred among European groups and *iii)* a weaker albeit significant regional clustering among American populations as previously observed (Verspoor 2005; Moore et al. 2014). These analyses, together with the clinal variation found by bourret et al. (2013) sets suitable conditions to reconstruct the underlying demographic processes that were involved in shaping the contemporary genetic landscape of Atlantic salmon throughout its natural distribution.

### Resolving the history of Atlantic salmon divergence

One of the most striking patterns of our demographic inferences was the unambiguous support for a scenario of long divergence without gene flow followed by very recent episodes of secondary contacts (approximately 1% of the total divergence time) in more than 90% of our comparisons, between continents, as well as among European and among North American populations. These long periods of isolation followed by relatively small periods of secondary contacts provided the best conditions to robustly infer secondary contacts (Roux et al. 2016). This is because long secondary contact periods result in the loss of isolation signal (Barton and Hewitt 1985) and that a situation of migration-drift equilibrium with a semi-permeable barrier to gene flow is attained (Endler 1977). This is often observed when reconstructing demography using only putatively “neutral” markers (Bierne et al. 2013; Rougemont et al. 2016).

We also observed a global reduction of genetic diversity among North American populations, as previously reported based on various molecular markers (Moore et al. 2014; Verspoor et al. 2005; King et al. 2007). Accordingly, *Treemix* analyses revealed higher levels of drift among North American populations relative to European ones. These results therefore support the hypothesis that North American populations were established by ancestral European populations (King et al. 2007). Indeed, similar patterns of reduced genetic diversity in newly founded populations relative to ancestral ones have commonly been reported, the most classical case being the “the out of Africa bottleneck(s) in humans” (Ramachandran et al. 2005; Henn et al. 2016).This continental divergence raises several questions: how long have the populations on each continent been isolated? When did the secondary contact start? Which were the sources of colonization of America? What is the potential impact of these small contemporary gene flow events on the genomic make up of contemporary populations?

We cannot provide direct absolute estimate of divergence time in years in our results section without formulating assumptions regarding generation time and size of reference (*N*ref as scaling factor in coalescent simulations. Nevertheless, using our best estimates of split time under a strict isolation model (n = 52 pairs of populations), assuming a mean generation time of four years (Palstra et al. 2009) and a *Nref* = 5, 000 (scaling factor in coalescent simulations) suggests that divergence between continents was initiated ~1,670,000 years ago [95% credible intervals =1,564,000−1,764,000], while the secondary contact between continental populations started ~ 13,400 years ago [95% credible intervals =1,800−41,380]. Our estimates of split times are therefore closer to those discussed by Nilsson et al. (2001)(>1 million years) than those of (King et al. 2007) (600 000 −700 000 years). These difference can be explained by the fact that the authors assumed different rate of substitution (1.2% per My (King et al. 2007) versus 0.5 to 0.9% per My to the same data Nilsson et al. 2001). Here, by estimating split time using the most highly differentiated loci according to their *F*_ST_ value, under a strict isolation model, we target loci that possess the deepest coalescence related to the onset of divergence. Our estimates suggested that continental divergence occurred during the mid-Pleistocene which lasted from ~ 2.58 Myrs ago to ~ 11,000 years ago (Gibbard et al. 2010). Accordingly, Atlantic Salmon on each side of the continent would have been in strict isolation approximately 99% of their divergence time during the Quaterny, a period where most of the earth surface was glaciated (Hewitt 2000). This long continental isolation period has most likely facilitated the accumulation of genetic incompatibility. This hypothesis is supported by the observation of pronounced asymmetric outbreeding depression at the second generation of hybridization reflecting the expression of Dobzhanskhy-Muller incompatibilities (Cauwelier et al. 2012). Thus, by crossing Canadian and Scottish salmon, the authors observed complete unviability of BC with Canadian fish whereas F1 and BC1 with Scottish fish were viable. Therefore Atlantic Salmon from each continent can be best described as partially reproductively isolated species, separated by a semi-permeable barrier to gene flow, as proposed recently in other systems such as European seabass *dicentrarchus labrax* (Tine et al. 2014); European Anchovy *Engraulis encrasicolus* (Le Moan et al. 2016) or European river and brook lampreys *Lampetra fluvatilis* and *L. planeri* (Rougemont et al. 2017). During the long glacial period secondary contacts were therefore unlikely.

Our results also suggested that following this long phase of geographic isolation, intercontinental secondary contact would have occurred at the end of the last glacial maxima (LGM), approximately 15 000 years ago. This period was associated with major environmental changes such as melting of ice sheets in North America and Europe, followed by an abrupt rise in sea levels as well as changes in oceanic circulations and temperature warming (Clark et al. 2009; Negre et al. 2010). These factors may have facilitated recent gene flow over long distance and impacted the current distribution and demographic history of species (Hewitt 1996; Bernatchez and Wilson 1998).

### Demographic history of European populations

A previous investigation (Bourret et al. 2013) found a cline in allele frequency at the majority of outlier loci when comparing the Baltic *versus* Atlantic populations and the Barents-White *versus* Atlantic populations, and the authors hypothesized that this reflected secondary contact between these genetic groups. Earlier studies have also proposed that Baltic populations must have existed in a separate refugium located either in the North Sea (Verspoor et al. 1999) or in the glacial lakes of Eastern Europe (Consuegra et al. 2002; King et al. 2007). More recently, a refugium was identified in Northern France with a potential contact zone between Northern France and the Iberian southern refugia (Finnegan et al. 2013). Here we validated the hypothesis of secondary contact around the Baltic Sea as well as in the Barents-White Sea. These results, together with these studies suggest that at least three refugia have existed (i.e. North Sea, Iberian Peninsula, Northwest France). We did not estimate divergence time under the secondary contact model as posteriors were not different from the priors and resulted in large uncertainty with regards to those estimates (i.e credible intervals = 239,000 – 2,230,000 years old). These results have important implications for interpreting genetic-environmental associations in this species. In particular, while endogenous barriers are most easily accumulated in allopatry and form tension zones upon secondary contact, their coupling with environmental barriers often stabilizes them (Barton 1979; Barton and Hewitt 1985) resulting in spurious genetic-environmental associations that may incorrectly be interpreted as local adaptation (Bierne et al. 2011). In the Baltic-Atlantic comparison, most of the differentiation observed in several species is generally attributed to adaptation to environmental gradients (e.g. Johannesson and André 2006; Gaggiotti et al. 2009; Berg et al. 2015; Guo et al. 2015)) under the hypothesis that the populations have adapted after the establishment of the Baltic sea (< 8000 years old). Here, our analysis indicates that this may not necessarily be the case. Without rejecting a potential secondary role of exogeneous barriers, our results show that the null model of demographic history can well account for the observed pattern, as observed in *Drosopgila. melanogaster*, for instance (Flatt 2016). We therefore propose that for any species, reconstructing the demographic history occurring along environmental gradients where hybrid zones have been described (e.g. Daguin et al. 2001; Bierne et al. 2003; Riginos and Cunningham 2005; Nikula et al. 2008) will allow constructing appropriate null model to better understand the relative role of demographic history *versus* environmental adaptation.

### Demographic history of North American populations

Our inferences of broad patterns of population genetic structure indicated two major groups with a north-south clustering. Among those, 350 out of 1095 (i.e. 32%) displayed mixed membership probabilities. This raises the same questions as for European populations, namely, does this pattern of contemporary admixture reflects divergence with gene flow or secondary contacts? First, the ABC analysis between the most differentiated groups provided strong support for secondary contacts. However, as for European populations, it was difficult to estimate the timing of divergence of the groups as credible intervals were large (22, 900 – 1,600,000years old). Second, we identified several European populations as source of admixture together with several major American sources from the southern area (Nova Scotia and Narraguagus, Québec lower and Upper Shore). This suggests that throughout its divergence history, the colonization of America by the ancestral European salmon populations has neither been established by a single colonization event nor by a single point of secondary contact. Interestingly, the Narraguagus River is the southernmost sample from our study and appears as the most extensive source of admixture in more northern populations. Admittedly however, the Narraguagus River may not be the actual source population itself but rather being representative of the southernmost range distribution of Atlantic Salmon, given that we could integrate only one population from USA. Therefore, it would be relevant to integrate more populations from this area.

Besides the strong signal of admixture from the southern region represented by Narraguagus River population, our results also suggest that several colonization events from different European populations into at least two major genetic groups occurred. The inference of admixture from two ancestral European branches into the Gaspésie and Anticosti areas further supports a hypothesis of ancestral, multiple colonization events. Therefore, we propose that North American populations most likely represent a mixture of multiple European lineages that varies among populations. As such, our results do not support the hypothesis of a single colonization event and single intercontinental secondary contact. Instead, a model of multiple colonization and contacts seem more plausible as was recently proposed to explain the colonization of North America by *D. melanogaster* (Brawand et al. 2014) in which North American populations would represent a mixture of European and African lineages. Also it may seem paradoxical to observe a lower genetic diversity of American population in spite of potential admixture. However, we argue that admixture resulting from different colonization event has mainly proceeded through a series of founding events and was more likely to be the results of a few individuals successfully migrating to North America. In scenario of a few individuals colonizing North America, drift certainly played a major role. Therefore, in spite of admixture, the full extent of ancestral genetic diversity is unlikely to be present in North American samples. Finally, given the potential strong incompatibilities of American and European genetic background, it is likely that the most recent (postglacial) introgression events are selected against and results in minor change in overall patterns of genetic diversity.

Moreover, our ABC modeling suggests that following these colonization events, spatial redistribution of European ancestral variants into neighboring American rivers apparently resulted in a complex signal of admixture integrating multiple signals of split and secondary contacts. Finally, as for Europe, results for North America indicate that studies focusing on interpretation of local adaptation may benefit from accounting for the possible confounding effects of admixture in generating allele frequency clines (Kapun et al. 2016). For instance, previous inferences of local adaptation in Atlantic Salmon in America have yield strong support for an association between spatial variation at MHC, temperature and pathogen diversity (Dionne et al. 2008) or between geological substrate and spatial genetic variation (Bourret et al. 2013b). It would be interesting to compare results of these previous studies with genome-scan based on our explicit demographic modeling.

### Evidence for genome wide heterogeneity and a role for linked selection

Another salient result of our study was that models incorporating heterogeneity of either Ne, *m* or both to account for linked selection outcompete models with homogeneous gene flow (except in three out of 148 cases). It is increasingly recognized that integrating heterogeneity of introgression rate (*m*) in demographic inferences increases the accuracy of model selection (Roux et al. 2013; Le Moan et al. 2016; Leroy et al. 2017; Rougemont et al. 2017) as genetic barriers to gene flow are known to reduce the effective rate of migration and to result in heterogeneous landscape of differentiation. A recent study (Roux et al. 2016) demonstrated the importance of integrating local genomic variation in *Ne* to model variation in intensities of genetic hitchhiking due to selective sweeps (Smith and Haigh 1974) or to background selection (Charlesworth et al. 1993). More specifically, these authors showed how neglecting one of these two components can lead to false inferences about intensities of gene flow, local variation in effective population size and ultimately impact model choice. Moreover, there is mounting evidence supporting a role of linked selection in shaping heterogeneous landscape of differentiation (Burri et al. 2015; Vijay et al. 2016; Vijay et al. 2017) and that gene flow is not always necessary to explain heterogeneous divergence. Here, however, our modeling approach suggests that gene flow associated with secondary contact has played an important role by lowering genome wide differentiation outside of barrier loci. Admittedly, our ABC framework is a simplified approach that does not allow distinguishing the process underlying local variation in *Ne*. It is likely that during the majority of the isolation period of Atlantic Salmon, background selection may have played a role, as suggested in other studies (Corbett-Detig et al. 2015). However, disentangling the relative contribution of differential introgression, selective sweeps and background selection was beyond the scope of this study and would require more data than those available so far in Atlantic Salmon, in particular with regards to the genome-wide estimates of recombination rates (e.g.Vijay et al. 2016). Given widespread variation in recombination rates in salmonids (Sakamoto et al. 2000; Moen et al. 2004; Sutherland et al. 2016) and evidence for a lower recombination near centromeres it would be relevant to perform an investigation across the salmonids radiation, as was recently done in birds (Vijay et al. 2017) in order to better understand if linked selection has also purged diversity over large evolutionary timescales.

## Conclusions

The modeling approach that we used here suggests that the demographic history of Atlantic Salmon divergence was shaped by multiple secondary contacts both between and within the North American and European continents. Multiple contact zones from European populations in North America, followed by widespread admixture and sorting of ancestral variation seems the most likely scenario for the observed patterns. While differential introgression across the genome certainly played a role in the formation of a heterogeneous differentiation pattern, results also clearly point to a role for linked selection in the form of background selection and/or positive hitchhiking. In these conditions, identifying targets of local adaptation will be particularly challenging. Clearly, more extensive genome wide data with information about local variation in recombination rate will be needed to address this issue (e.g. centromeres, inversions). These data will also allow drawing models of divergence with more than two populations and will allow using more appropriate null neutral model for detecting targets of selection associated with local adaptation. With all these confounding effects in mind, disentangling real targets of adaptation from demographic and other nonadaptive processes may be more challenging than has been appreciated thus far.

## Methods

### Sampling and genotyping

A total of 2,080 individuals from 77 locations across the whole native range of Atlantic Salmon from two studies (Bourret et al. 2013, Moore et al, 2014)(Fig. 1) were merged which resulted in a total of 5,087 loci from a SNP array previously developed at the Centre for Integrative Genetics (CIGENE) in Norway. All details from the SNP discovery as well as quality control and ascertainment bias in North American populations are described in Bourret et al. (2013). Genotypes were further filtered by excluding individuals with more than 5% missing genotypes (45 individuals removed) as well as SNPs with a lower than 95% genotyping rate (53 SNPs removed). Only SNPs with a global minor allele frequency (MAF) > 5% were kept (379 SNPs removed) resulting in a final dataset of 2,035 individuals and 4656 SNPs with less than 1% of missing data. Basic summary statistics (i.e. Ho, He, pairwise *F*_ST_ were computed globally and for each locus using mainly custom R scripts (see Data Availability section) and Hiersfstat package (Goudet, 2005).

### Patterns of broad population genetic structure and admixture

We first visualized population genetic relationships among rivers and individuals using a principal component analysis as implemented in the R package ade4 (Dray and Dufour 2007) using the whole dataset. Second, we inferred levels of ancestry and admixture proportion of individuals using the *snmf* function implemented in the R package LEA (Frichot et al. 2015). K-values ranging between one and 60 were tested and cross-validations were obtained using the cross-entropy criterion with 5% of masked genotypes. The default value for the regularization parameter was used to avoid forcing individuals into groups hence underestimating admixture. Maps of ancestry coefficients were drawn in R following recommendations available elsewhere (Caye et al. 2016).

### SNP mapping on the Atlantic salmon reference genome

The *Treemix* analysis is best performed when SNPs are positioned in the genome. Therefore, the recently published Atlantic Salmon genome was used (accession code: PRJNA72713 and PRJNA260929 NCBI) to map SNPs on the genome (Lien et al. 2016). The masked reference genome was used to avoid mapping on repeat sequences. Reads containing SNPs were mapped on the genome using gsnap (Wu and Nacu 2010) with default parameters. Reads were then converted to bam files using samtools (Li et al. 2009) and unmapped or secondary mapped reads were removed. A total of 4,637 of 4,656 (99%) reads were successfully mapped on the genome. Only high quality mappings were retained (MAPQ>40) resulting in a total of 4153 SNPs. We also exploited these newly positioned SNPs to plots overall *F*_ST_ values along each chromosomes (Supplemental Fig S1).

## Demographic inference

### Inferring population splits and sources of admixture

We combined TreeMix v1.12 (Pickrell and Pritchard, 2012), *f3-test* (Reich et al. 2009) and Spacemix (Bradburg et al. 2016) to identify population splits and admixture as well as the most likely source(s) of admixture. Treemix uses allele frequency covariance matrix and models genetic drift through a Gaussian approximation to build the maximum likelihood population graph. Migration events were set between 0 and 20 aiming to reach an asymptotic value in the percentage of variance explained (Supplemental Fig S6).

Second, we used the *f3-s*tatistical tests (Reich et al. 2009) with bootstraps in blocks of 100 SNPs to test whether divergence between European and North American populations can be best explained by a model without migration. The *f3*-statistical tests or three-population test is a formal test of admixture (f3(C; A, B)) which tests whether population C derived a significant amount of admixture from population A and B. All possible comparisons of three populations were performed.

Finally, we used a spatial framework for inference of the geogenetic positions of populations with their most probable source and direction of admixture implemented in the R package SpaceMix (Bradburg et al. 2016). The goal of this analysis was to complement Treemix analyses by using an explicit geographic framework in order to identify the potential sources of admixture and directions of migration events. Two demographic models were tested. In the first model, only population locations were estimated but not admixture, corresponding to a model of pure isolation by distance. The second model estimated together the population locations as well as the extent of admixture (see details in (Bradburg et al. 2016)). Parameter settings strictly followed those implemented by the authors for the study of human populations. Analyses were performed using the covariance-matrix on the full dataset. The convergence of the runs was checked using functions provided in the SpaceMix package

### Explicit modeling: coalescent analysis and Approximate Bayesian Computation

A new ABC framework recently developed (Roux et al. 2013, 2016) was used to perform explicit modeling of divergence history while incorporating the effects of selection at linked sites affecting *Ne* and differential introgression (*m*).

### Coalescent analyses

Demographic scenario were tested on a subset of populations representative of the major genetic groups identified in Bourret et al. (2013) for Europe, Moore et al. (2014) for North America and here. The list of all major groups is provided in S1 table. A total of 15 groups were defined and pairwise comparison of divergence scenarios was performed as detailed below.

Four classic demographic scenarios were compared (Fig. 1). These include a model of strict isolation (SI), a model of divergence with migration during the first generations of divergence or ancestral migration (AM), a model of divergence with continuous gene flow or isolation with migration (IM) and a model of secondary contact (SC). Each model is characterized by a set of demographic parameters and assumptions. The SI model assumes an instantaneous split at *T_split_* of an ancestral population of size *N_ANC_* into two daughter populations of constant size *N*_pop1_ and *N*_pop2_ and no gene flow. The AM model assumes an instantaneous split of the ancestral population followed by a period of gene flow from *T_split_* to *T_am_* and then a period without gene flow until the present date. The IM model assumes that after *T_split_*, the two daughter populations continuously exchange gene at a constant rate each generation. The SC model assumes an initial period of strict isolation without gene flow after *T_split_* and is followed by a period of secondary contact *T*_sc_ generations ago that is still ongoing. The IM, AM and SC models included migration as *M* = 4 *N_0_.m*, with M_1←2_being the number of migrants from population 2 to population 1 and M2←1being the reciprocal. Then heterogeneity of migration and of effective population size was modeled using beta distributions as hyper-prior on each of these two parameters (see below). For each model, 1×10^6^ simulations of datasets matching the sample size of each locus in each pair-wise dataset was performed using msnsam (Ross-Ibarra et al. 2008) a modified version of the coalescent simulator ms (Hudson 2002) under an infinite-site mutation model. Large and uninformative priors were drawn as follows: priors for the divergence time (*T*_split_/4*Ne*) were uniformly sampled on the interval [0–30], so that *T*_am_, *T*_sc_ and *T*_si_ were constrained to be chosen within this interval. For homogeneous *Ne, N_ANC_ N_Pop1_*,and *N_Pop2_* were sampled independently and uniformly on the interval [0–20]. For the homogeneous version of the migration rate, *M* was sampled on the interval [0–40] independently for each direction of migrations. For model including genomic heterogeneity in introgression rate heterogeneity was modeled using a beta distribution as hyperprior shaped by two parameters; α randomly sampled on the interval [0–20] and β randomly sampled on the interval [0–200]. A value was then independently assigned to each locus. For heterogeneity of *Ne*, α percent of loci shared a common value uniformly sampled on the interval [0–1] and represents the proportion of loci assumed to evolve neutrally. Then a proportion 1 − α was assumed to be under the effect of selection at linked sites. Their *Ne* values were thus drawn from a beta distribution defined on the interval [0–20], as defined on the homogeneous version. *Ne* values were thus independently drawn for *N*_ANC_, *N*_1_ and *N*_2_ but shared the same shape parameters (Roux et al. 2016). Priors were generated using a Python version of priorgen software (Ross-Ibarra et al. 2008) and a panel of commonly used summary statistics (Fagundes et al. 2007; Roux et al. 2013) was computed using mscalc (Ross-Ibarra et al. 2008; Ross-Ibarra et al. 2009; Roux et al. 2011).

### Integrating multiple episodes of gene-flow and demographic size change

From the best inferred model, (systematically SC, see results), we tested whether integrating two events of secondary contact and two events of strict isolation under a model called periodic secondary contact (PSC) best fitted our data than the model of single secondary contact (Merceron et al. 2017). This model was designed to better reflect the possible outcome of multiple glacial expansions and retreats throughout historical times. Similarly, another expected outcome of divergence and gene flow during post-glacial colonization is that the colonization of neighboring rivers by few founding individuals may generate strong drift and result in bottleneck in the newly founded populations (BSC). To test these two hypotheses, we chose a completely random subset of population pairs derived from the 162 pairs analyzed previously. In order to be representative of the previous analyses, we included pairs diverging between continents, between major genetic groups within Europe and within America. We thus modified our inference pipeline to explore the fit of these models to our data. Details of the modifications are provided in Supplemental Note S1.

### ABC: model selection

A total of 14 models were available: four alternative models for IM, AM, SC hereafter called *NhomoMhomo* (for Homogeneous *Ne* and Homogeneous migration along the genome); *NhomoMhetero* (for Homogeneous *Ne* and Heterogeneous migration along the genome); *NheteroMhomo* (for Heterogeneous *Ne* and heterogeneous migration along the genome) and *NheteroMhetero* (for Heterogeneous *Ne* and heterogeneous migration along the genome); and two alternative models for SI that is *Nhomo* or *Nhetero*. Based on the identification of the best model, we further constructed two additional models incorporating periodic secondary contact (PSC) and variation in effective population size (BSC).

First, the 14 models were all compared together at the same time using ABC. Posterior probabilities were computed by retaining the 3500 ‘best’ simulations (out of 14 × 1 million) during the rejection step, those were weighted by an Epanechnikov kernel peaking when *S*_obs_ = *S*_sim_and a neural network was used for the rejection step. Fifty neural networks were used for training with 15 hidden layers. Ten replicates were performed to compute an average posterior probability for each model. Last, we compared the best secondary contact models against the two remaining alternatives PSC and BSC. Each time, the same model selection procedure was repeated. The R package 'abc' (Csilléry et al. 2012) was used for the model choice procedure.

### Robustness

The robustness, which is defined as the probability of correctly classifying a model given a posterior probability threshold was assessed using pseudo-observed dataset (PODS). A total of 10,000 PODS from each model was simulated with parameters drawn from the same prior distribution. Then, the same ABC model choice procedure as above was run again, but using the pseudo-observed dataset instead of the real data set in order to compute the posterior probability of each PODS. From the posterior probability distribution the robustness was then computed as defined elsewehere (eg: Roux et al. (2016)).

### Parameter estimation and posterior predictive checks

For every robustly inferred model, parameter estimation was performed using a logit transformation of the parameters and a tolerance of 0.001. The posterior probabilities of parameter values were then estimated using the neural network procedure with nonlinear regressions of the parameters on the summary statistics using 50 feed-forwards neural networks and 15 hidden layers. We then performed posterior predictive check (ppc) to assess the goodness of fit of the best supported model (Gelman 2003; Rougemont et al. 2016). We ran 2,000 simulations for each locus using the joint posterior parameter estimates in each comparison to generate summary statistics and compare them to those empirically observed in each dataset. The simulation pipeline was used again to do so and to compute the p-value for each summary statistic in order to estimate the fit.

#### Data availability

The whole pipeline used to perform demographic inference, is available at: <u>https://github.com/QuentinRougemont/abc_inferences</u>

## Acknowledgements

We are grateful to Eric Normandeau for his help in setting up some of the bioinformatics pipelines implemented in this study. We are deeply indebted to Thibault Leroy and Camille Roux for great discussions around ABC inferences. Thank you also to Clément Rougeux, Anne-Laure Ferchaud, Ben Sutherland and JS Moore for their comments on an earlier version of the manuscript. This study was funded by the Canada Research Chair in Genomics and Conservation of Aquatic Resources and an NSERC (Canada) Discovery grant held by LB.

## References

Barton N, Bengtsson BO. 1986. The barrier to genetic exchange between hybridising populations. Heredity 57:357–376.

Barton NH. 1979. The dynamics of hybrid zones. Heredity 43:341–359.

Barton NH, Hewitt GM. 1985. Analysis of Hybrid Zones. Annu. Rev. Ecol. Syst. 16:113–148.

Beaumont MA, Zhang W, Balding DJ. 2002. Approximate Bayesian Computation in Population Genetics. Genetics 162:2025–2035.

Berg PR, Jentoft S, Star B, Ring KH, Knutsen H, Lien S, Jakobsen KS, Andre C. 2015. Adaptation to Low Salinity Promotes Genomic Divergence in Atlantic Cod (*Gadus morhua L.*). Genome Biol. Evol. 7:1644–1663.

Bernatchez L, Wilson CC. 1998. Comparative phylogeography of Nearctic and Palearctic fishes. Mol. Ecol. 7:431–452.

Bierne N, Borsa P, Daguin C, Jollivet D, Viard F, Bonhomme F, David P. 2003. Introgression patterns in the mosaic hybrid zone between Mytilus edulis and M. galloprovincialis. Mol. Ecol. 12:447–461.

Bierne N, Gagnaire P-A, David P. 2013. The geography of introgression in a patchy environment and the thorn in the side of ecological speciation. Cur. Zool. 59: 72—86.

Bierne N, Welch J, Loire E, Bonhomme F, David P. 2011. The coupling hypothesis: why genome scans may fail to map local adaptation genes. Mol. Ecol. 20:2044–2072.

Bolstad GH, Hindar K, Robertsen G, Jonsson B, Sægrov H, Diserud OH, Fiske P, Jensen AJ, Urdal K, Næsje TF, et al. 2017. Gene flow from domesticated escapes alters the life history of wild Atlantic salmon. Nat. Ecol. Evol. 1:0124.

Bourke E. 1997. Allozyme variation in populations of Atlantic salmon located throughout Europe: diversity that could be compromised by introductions of reared fish. ICES J. Mar. Sci. 54:974–985.

Bourret V, Kent MP, Primmer CR, Vasemägi A, Karlsson S, Hindar K, McGinnity P, Verspoor E, Bernatchez L, Lien S. 2013. SNP-array reveals genome-wide patterns of geographical and potential adaptive divergence across the natural range of Atlantic salmon (*Salmo salar*). Mol. Ecol. 22:532–551.

Bourret V, Dionne M, Kent MP, Lien S, Bernatchez L. 2013. Landscape genomics in Atlantic Salmon (*Salmo salar*): Searching for environment interactions driving local adaptation. Evolution 67:3469–3487.

Bradburd GS, Ralph PL, Coop GM. 2016. A Spatial Framework for Understanding Population Structure and Admixture. Slatkin M, editor. PLOS Genet. 12:e1005703.

Bradbury IR, Hamilton LC, Dempson B, Robertson MJ, Bourret V, Bernatchez L, Verspoor E. 2015. Transatlantic secondary contact in Atlantic Salmon, comparing microsatellites, a single nucleotide polymorphism array and restriction-site associated DNA sequencing for the resolution of complex spatial structure. Mol. Ecol. 24:5130–5144.

Brawand D, Wagner CE, Li YI, Malinsky M, Keller I, Fan S, Simakov O, Ng AY, Lim ZW, Bezault E, et al. 2014. The genomic substrate for adaptive radiation in African cichlid fish. Nature 513:375–381.

Burri R, Nater A, Kawakami T, Mugal CF, Olason PI, Smeds L, Suh A, Dutoit L,

Bureš S, Garamszegi LZ, et al. 2015. Linked selection and recombination rate variation drive the evolution of the genomic landscape of differentiation across the speciation continuum of Ficedula flycatchers. Genome Res. 25:1656–1665.

Cauwelier E, Gilbey J, Jones CS, Noble LR, Verspoor E. 2012. Asymmetrical viability in backcrosses between highly divergent populations of Atlantic salmon (Salmo salar): implications for conservation. Conserv. Genet. 13:1665–1669.

Caye K, Deist TM, Martins H, Michel O, François O. 2016. TESS3: fast inference of spatial population structure and genome scans for selection. Mol. Ecol. Resour. 16:540–548.

Charlesworth B. 1994. The effect of background selection against deleterious mutations on weakly selected, linked variants. Genet. Res. 63:213–227.

Charlesworth B, Morgan MT, Charlesworth D. 1993. The effect of deleterious mutations on neutral molecular variation. Genetics 134:1289–1303.

Clark PU, Dyke AS, Shakun JD, Carlson AE, Clark J, Wohlfarth B, Mitrovica JX, Hostetler SW, McCabe AM. 2009. The Last Glacial Maximum. Science 325:710–714.

Consuegra S, García De Leániz C, Serdio A, González Morales M, Straus LG, Knox D, Verspoor E. 2002. Mitochondrial DNA variation in Pleistocene and modern Atlantic salmon from the Iberian glacial refugium. Mol. Ecol. 11:2037–2048.

Corbett-Detig RB, Hartl DL, Sackton TB. 2015. Natural Selection Constrains Neutral Diversity across A Wide Range of Species. Barton NH, editor. PLOS Biol. 13:e1002112.

Cruickshank TE, Hahn MW. 2014. Reanalysis suggests that genomic islands of speciation are due to reduced diversity, not reduced gene flow. Mol. Ecol. 23:3133–3157.

Csilléry K, François O, Blum MGB. 2012. abc: an R package for approximate Bayesian computation (ABC). Methods Ecol. Evol. 3:475–479.

Cutler MG, Bartlett SE, Hartley SE, Davidson WS. 1991. A Polymorphism in the Ribosomal RNA Genes Distinguishes Atlantic Salmon (*Salmo salar*) from North America and Europe. Can. J. Fish. Aquat. Sci. 48:1655–1661.

Daguin C, Bonhomme F, Borsa P. 2001. The zone of sympatry and hybridization of Mytilus edulis and M. galloprovincialis, as described by intron length polymorphism at locus mac-1. Heredity 86:342–354.

Dionne M, Caron F, Dodson JJ, Bernatchez L. 2008. Landscape genetics and hierarchical genetic structure in Atlantic salmon: the interaction of gene flow and local adaptation: Landscape genetics in Atlantic Salmon. Mol. Ecol. 17:2382–2396.

Dray S, Dufour A-B. 2007. The ade4 Package: Implementing the Duality Diagram for Ecologists. J. Stat. Softw. [Internet] 22. Available from: http://www.jstatsoft.org/v22/i04/

Endler JA. 1977. Geographic Variation, Speciation, and Clines. Princeton University Press

Ewing GB, Jensen JD. 2016. The consequences of not accounting for background selection in demographic inference. Mol. Ecol. 25:135–141.

Fagundes NJR, Ray N, Beaumont M, Neuenschwander S, Salzano FM, Bonatto SL, Excoffier L. 2007. Statistical evaluation of alternative models of human evolution. Proc. Natl. Acad. Sci. 104:17614–17619.

Falush D, Dorp L van, Lawson D. 2016. A tutorial on how (not) to over-interpret STRUCTURE/ADMIXTURE bar plots. bioRxiv:066431. https://doi.org/10.1101/066431, last accessed September 2016.

Feder JL, Egan SP, Nosil P. 2012. The genomics of speciation-with-gene-flow. Trends Genet. 28:342–350.

Ferchaud A-L, Perrier C, April J, Hernandez C, Dionne M, Bernatchez L. 2016. Making sense of the relationships between Ne, Nb and Nc towards defining conservation thresholds in Atlantic salmon (*Salmo salar*). Heredity 117:268–278.

Finnegan AK, Griffiths AM, King RA, Machado-Schiaffino G, Porcher J-P, Garcia-Vazquez E, Bright D, Stevens JR. 2013. Use of multiple markers demonstrates a cryptic western refugium and postglacial colonisation routes of Atlantic salmon (*Salmo salar L.*) in northwest Europe. Heredity 111:34–43.

Flatt T. 2016. Genomics of clinal variation in *Drosophila*: disentangling the interactions of selection and demography. Mol. Ecol. 25:1023–1026.

Flaxman SM, Feder JL, Nosil P. 2013. Genetic Hitchhiking and the Dynamic Buildup of Genomic Divergence During Speciation with Gene Flow. Evolution 67:2577–2591.

Frichot E, François O. 2015. LEA: An R package for landscape and ecological association studies. O’Meara B, editor. Methods Ecol. Evol. 6:925–929.

Gaggiotti OE, Bekkevold D, Jørgensen HBH, Foll M, Carvalho GR, Andre C, Ruzzante DE. 2009. Disentangling the Effects of Evolutionary, Demographic, and Environmental Factors Influencing Genetic Structure of Natural Populations: Atlantic Herring as a Case Study. Evolution 63:2939–2951.

Gagnaire P-A, Pavey SA, Normandeau E, Bernatchez L. 2013. The Genetic Architecture of Reproductive Isolation During Speciation-with-Gene-Flow in Lake Whitefish Species Pairs Assessed by Rad Sequencing. Evolution 67:2483–2497.

Gelman A. 2003. A Bayesian Formulation of Exploratory Data Analysis and Goodness-of-fit Testing*. Int. Stat. Rev. 71:369–382.

Gibbard PL, Head MJ, Walker MJC, the Subcommission on Quaternary Stratigraphy. 2010. Formal ratification of the Quaternary System/Period and the Pleistocene Series/Epoch with a base at 2.58 Ma. J. Quat. Sci. 25:96–102.

Guo B, DeFaveri J, Sotelo G, Nair A, Merilä J. 2015. Population genomic evidence for adaptive differentiation in Baltic Sea three-spined sticklebacks. BMC Biol. [Internet] 13. Available from: http://www.biomedcentral.com/1741-7007/13/19

Harrison RG, Larson EL. 2016. Heterogeneous genome divergence, differential introgression, and the origin and structure of hybrid zones. Mol. Ecol. 25:2454–2466.

Hartley SE. 1987. The chromosomes of Salmonid Fishes. Biol. Rev. 62:197–214.

Henn BM, Botigué LR, Peischl S, Dupanloup I, Lipatov M, Maples BK, Martin AR, Musharoff S, Cann H, Snyder MP, et al. 2016. Distance from sub-Saharan Africa predicts mutational load in diverse human genomes. Proc. Natl. Acad. Sci. 113:E440–E449.

Hewitt G. 2000. The genetic legacy of the Quaternary ice ages. Nature 405:907–913.

Hewitt GM. 1996. Some genetic consequences of ice ages, and their role in divergence and speciation. Biol. J. Linn. Soc. 58:247–276.

Hill WG, Robertson A. 1966. The effect of linkage on limits to artificial selection. Genet. Res. 8:269–294.

Hudson RR. 2002. Generating samples under a Wright-Fisher neutral model of genetic variation. Bioinforma. Oxf. Engl. 18:337–338.

Johannesson K, André C. 2006. Life on the margin: genetic isolation and diversity loss in a peripheral marine ecosystem, the Baltic Sea:Genetics of marginal Baltic Sea populations. Mol. Ecol. 15:2013–2029.

Kapun M, Fabian DK, Goudet J, Flatt T. 2016. Genomic Evidence for Adaptive Inversion Clines in *Drosophila melanogaster*. Mol. Biol. Evol. 33:1317–1336.

King TL, Verspoor E, Spidle AP, Gross R, Phillips RB, Koljonen M-L, Sanchez JA, Morrison CL. 2007. Biodiversity and Population Structure. In: Verspoor Eric, Stradmeyer L, Nielsen J, editors. The Atlantic Salmon. Oxford, UK: Blackwell Publishing Ltd. p. 117–166. Available from: http://doi.wiley.com/10.1002/9780470995846.ch5

Le Moan A, Gagnaire P-A, Bonhomme F. 2016. Parallel genetic divergence among coastal-marine ecotype pairs of European anchovy explained by differential introgression after secondary contact. Mol. Ecol. 25:3187–3202.

Leroy T, Roux C, Villate L, Bodénès C, Romiguier J, Paiva JAP, Dossat C, Aury J-M, Plomion C, Kremer A. 2017. Extensive recent secondary contacts between four European white oak species. New Phytol.:n/a-n/a.

Li H, Handsaker B, Wysoker A, Fennell T, Ruan J, Homer N, Marth G, Abecasis G, Durbin R, 1000 Genome Project Data Processing Subgroup. 2009. The Sequence Alignment/Map format and SAMtools. Bioinformatics 25:2078–2079.

Lien S, Koop BF, Sandve SR, Miller JR, Kent MP, Nome T, Hvidsten TR, Leong JS, Minkley DR, Zimin A, et al. 2016. The Atlantic salmon genome provides insights into rediploidization. Nature 533:200–205.

Merceron NR, Leroy T, Chancerel E, Romero-Severson J, Borkowski D, Ducousso A, Monty A, Porté AJ, Kremer A. Back to America: tracking the origin of European introduced populations of Quercus rubra L. Genome. Forthcoming.

Milot E, Perrier C, Papillon L, Dodson JJ, Bernatchez L. 2013. Reduced fitness of Atlantic salmon released in the wild after one generation of captive breeding. Evol. Appl. 6:472–485.

Moen T, Hoyheim B, Munck H, Gomez-Raya L. 2004. A linkage map of Atlantic salmon (*Salmo salar*) reveals an uncommonly large difference in recombination rate between the sexes. Anim. Genet. 35:81–92.

Moore J-S, Bourret V, Dionne M, Bradbury I, O’Reilly P, Kent M, Chaput G, Bernatchez L. 2014. Conservation genomics of anadromous Atlantic salmon across its North American range: outlier loci identify the same patterns of population structure as neutral loci. Mol. Ecol. 23:5680–5697.

Negre C, Zahn R, Thomas AL, Masqué P, Henderson GM, Martínez-Méndez G, Hall IR, Mas JL. 2010. Reversed flow of Atlantic deep water during the Last Glacial Maximum. Nature 468:84–88.

Nikula R, Strelkov P, VäInöLä R. 2008. A broad transition zone between an inner Baltic hybrid swarm and a pure North Sea subspecies of Macoma balthica (Mollusca, Bivalvia). Mol. Ecol. 17:1505–1522.

Nilsson J, Gross R, Asplund T, Dove O, Jansson H, Kelloniemi J, Kohlmann K, LOytynoja A, Nielsen EE, Paaver T, et al. 2001. Matrilinear phylogeography of Atlantic salmon (*Salmo salar L.*) in Europe and postglacial colonization of the Baltic Sea area. Mol. Ecol. 10:89–102.

Noor M a. F, Bennett SM. 2009. Islands of speciation or mirages in the desert? Examining the role of restricted recombination in maintaining species. Heredity 103:439–444.

Palstra FP, O’connell MF, Ruzzante DE. 2007. Population structure and gene flow reversals in Atlantic salmon (*Salmo salar*) over contemporary and longterm temporal scales: effects of population size and life history. Mol. Ecol. 16:4504–4522.

Palstra FP, O’Connell MF, Ruzzante DE. 2009. Age Structure, Changing Demography and Effective Population Size in Atlantic Salmon (*Salmo salar*). Genetics 182:1233–1249.

Perrier C, Baglinière J-L, Evanno G. 2013. Understanding admixture patterns in supplemented populations: a case study combining molecular analyses and temporally explicit simulations in Atlantic salmon. Evol. Appl. 6:218–230.

Perrier C, Guyomard R, Bagliniere J-L, Evanno G. 2011. Determinants of hierarchical genetic structure in Atlantic salmon populations: environmental factors vs. anthropogenic influences. Mol. Ecol. 20:4231–4245.

Pickrell JK, Pritchard JK. 2012. Inference of Population Splits and Mixtures from Genome-Wide Allele Frequency Data. Tang H, editor. PLoS Genet. 8:e1002967.

Primmer CR. 2011. Genetics of local adaptation in salmonid fishes. Heredity 106:401–403.

Quinn TP. 1993. A review of homing and straying of wild and hatchery-produced salmon. Fish. Res. 18:29–44.

Ramachandran S, Deshpande O, Roseman CC, Rosenberg NA, Feldman MW, Cavalli-Sforza LL. 2005. Support from the relationship of genetic and geographic distance in human populations for a serial founder effect originating in Africa. Proc. Natl. Acad. Sci. 102:15942–15947.

Reich D, Thangaraj K, Patterson N, Price AL, Singh L. 2009. Reconstructing Indian population history. Nature 461:489–494.

Riginos C, Cunningham CW. 2005. Local adaptation and species segregation in two mussel (*Mytilus edulis x Mytilus trossulus) hybrid zones*. Mol. Ecol. 14:381–400.

Ross-Ibarra J, Tenaillon M, Gaut BS. 2009. Historical Divergence and Gene Flow in the Genus *Zea*. Genetics 181:1399–1413.

Ross-Ibarra J, Wright SI, Foxe JP, Kawabe A, DeRose-Wilson L, Gos G, Charlesworth D, Gaut BS. 2008. Patterns of Polymorphism and Demographic History in Natural Populations of *Arabidopsis lyrata*. Fay JC, editor. PLoS ONE 3:e2411.

Rougemont Q, Gagnaire P-A, Perrier C, Genthon C, Besnard A-L, Launey S, Evanno G. 2017. Inferring the demographic history underlying parallel genomic divergence among pairs of parasitic and nonparasitic lamprey ecotypes. Mol. Ecol. 26:142–162.

Rougemont Q, Roux C, Neuenschwander S, Goudet J, Launey S, Evanno G. 2016. Reconstructing the demographic history of divergence between European river and brook lampreys using approximate Bayesian computations. PeerJ 4:e1910.

Rougeux C, Bernatchez L, Gagnaire P-A. 2016. Modeling the multiple facets of speciation-with-gene-flow towards improving divergence history inference of a recent fish adaptive radiation. GBE. Forthcoming.

Roux C, Castric V, Pauwels M, Wright SI, Saumitou-Laprade P, Vekemans X. 2011. Does Speciation between Arabidopsis halleri and Arabidopsis lyrata Coincide with Major Changes in a Molecular Target of Adaptation? Ingvarsson PK, editor. PLoS ONE 6:e26872.

Roux C, Fraïsse C, Romiguier J, Anciaux Y, Galtier N, Bierne N. 2016. Shedding Light on the Grey Zone of Speciation along a Continuum of Genomic Divergence. Moritz C, editor. PLOS Biol. 14:e2000234.

Roux C, Tsagkogeorga G, Bierne N, Galtier N. 2013. Crossing the Species Barrier: Genomic Hotspots of Introgression between Two Highly Divergent Ciona intestinalis Species. Mol. Biol. Evol. 30:1574–1587.

Sakamoto T, Danzmann RG, Gharbi K, Howard P, Ozaki A, Khoo SK, Woram RA, Okamoto N, Ferguson MM, Holm LE, et al. 2000. A microsatellite linkage map of rainbow trout (*Oncorhynchus mykiss*) characterized by large sex-specific differences in recombination rates. Genetics 155:1331–1345.

Schrider DR, Shanku AG, Kern AD. 2016. Effects of Linked Selective Sweeps on Demographic Inference and Model Selection. Genetics 204:1207–1223.

Seehausen O, Butlin RK, Keller I, Wagner CE, Boughman JW, Hohenlohe PA, Peichel CL, Saetre G-P, Bank C, Brännström Å, et al. 2014. Genomics and the origin of species. Nat. Rev. Genet. 15:176–192.

Smith JM, Haigh J. 1974. The hitch-hiking effect of a favourable gene. Genet. Res. 23:23–35.

Sousa VC, Carneiro M, Ferrand N, Hey J. 2013. Identifying Loci Under Selection Against Gene Flow in Isolation-with-Migration Models. Genetics 194:211–233.

Sutherland BJG, Gosselin T, Normandeau E, Lamothe M, Isabel N, Audet C, Bernatchez L. 2016. Salmonid chromosome evolution as revealed by a novel method for comparing RADseq linkage maps. Genome Biol. Evol.:evw262.

Tavaré S, Balding DJ, Griffiths RC, Donnelly P. 1997. Inferring Coalescence Times From DNA Sequence Data. Genetics 145:505–518.

Taylor EB. 1991. A review of local adaptation in Salmonidac, with particular reference to Pacific and Atlantic salmon. Aquaculture 98:185–207.

Tine M, Kuhl H, Gagnaire P-A, Louro B, Desmarais E, Martins RST, Hecht J, Knaust F, Belkhir K, Klages S, et al. 2014. European sea bass genome and its variation provide insights into adaptation to euryhalinity and speciation. Nat. Commun. [Internet] 5. Available from: http://www.nature.com/ncomms/2014/141223/ncomms6770/full/ncomms6770.html

Vasemägi A, Nilsson J, Primmer CR. 2005. Expressed sequence tag-linked microsatellites as a source of gene-associated polymorphisms for detecting signatures of divergent selection in atlantic salmon (*Salmo salar L.*). Mol. Biol. Evol. 22:1067–1076.

Verspoor E. 2005. Regional differentiation of North American Atlantic salmon at allozyme loci. J. Fish Biol. 67:80–103.

Verspoor E, Beardmore JA, Consuegra S, García de Leániz C, Hindar K, Jordan WC, Koljonen M-L, Mahkrov AA, Paaver T, Sánchez JA, et al. 2005. Population structure in the Atlantic salmon: insights from 40 years of research into genetic protein variation. J. Fish Biol. 67:3–54.

Verspoor E, McCARTHY EM, Knox D, Bourke EA, Cross TF. 1999. The phylogeography of European Atlantic salmon (*Salmo salar L.*) based on RFLP analysis of the ND1/16sRNA region of the mtDNA. Biol. J. Linn. Soc. 68:129–146.

Via S, West J. 2008. The genetic mosaic suggests a new role for hitchhiking in ecological speciation. Mol. Ecol. 17:4334–4345.

Vijay N, Bossu CM, Poelstra JW, Weissensteiner MH, Suh A, Kryukov AP, Wolf JBW. 2016. Evolution of heterogeneous genome differentiation across multiple contact zones in a crow species complex. Nat. Commun. 7:13195.

Vijay N, Weissensteiner M, Burri R, Kawakami T, Ellegren H, Wolf JBW. 2017. Genome-wide signatures of genetic variation within and between populations - a comparative perspective. bioRxiv:104604. https://doi.org/10.1101/104604, Last accessed April 2017

Wolf JBW, Ellegren H. 2016. Making sense of genomic islands of differentiation in light of speciation. Nat. Rev. Genet. 18:87–100.

Wu C-I. 2001. The genic view of the process of speciation. J. Evol. Biol. 14:851–865.

Wu TD, Nacu S. 2010. Fast and SNP-tolerant detection of complex variants and splicing in short reads. Bioinformatics 26:873–881.

